# Chaperone-mediated autophagy is an overlooked pathway for mutant α1-antitrypsin Z degradation

**DOI:** 10.1101/2023.11.24.568525

**Authors:** Jiayu Lin, Haorui Lu, Xinyue Wei, Yan Dai, Rihan Wu, Hao Yang, Lang Rao

## Abstract

Chaperone-mediated autophagy (CMA) is a specific form of autophagy that selectively targets proteins containing a KFERQ-like motif and relies on the chaperone protein HSC70 for substrate recognition. In α1-antitrypsin deficiency (AATD), a disease characterized by the hepatic build-up of α1-Antitrypsin Z mutant (ATZ), CMA’s role had been unclear. This work demonstrates the critical role that CMA plays in preventing ATZ accumulation; suppressing CMA worsens ATZ accumulation, whilst activating it through chemical stimulation or LAMP2A overexpression promotes ATZ breakdown. Specifically, ATZ’s 121QELLR125 motif is critical for HSC70 recognition and LAMP2A’s charged C-terminal cytoplasmic tail is vital for substrate binding, facilitating CMA-mediated degradation of ATZ. This selective activation of CMA operates independently from other autophagy pathways and alleviate ATZ aggregates caused cellular stress. These findings highlight CMA’s critical function in cellular protein quality control of ATZ and place it as a novel target for AATD treatment approaches.

## Introduction

The autophagy-lysosome pathway is a major protein aggregate clearance mechanism that is conserved in all eukaryotic cells. This system can be classified into three main types based on their distinctive digestion processes: macroautophagy, microautophagy, and chaperone-mediated autophagy (CMA)[<u>1</u>]. Macroautophagy (referred to as autophagy) and microautophagy non-selectively engulf various substrates, whereas CMA specifically targets proteins containing the CMA-targeting KFERQ motif [<u>2,3</u>]. In CMA pathway, the heat shock cognate protein of 70 kDa (HSC70, also known as HSPA8) recognizes the KFERQ motif of substrate to form a substrate–chaperone complex [<u>4</u>], the complex was then delivered to lysosome surface to interact with the CMA specific receptor lysosome-associated membrane protein type 2A (LAMP2A). The LAMP2A in charge of delivering substrates directly to the lysosomal lumen for degradation [<u>4,5</u>]. Increasing evidence suggests that CMA plays a crucial role in various of physiological processes; CMA malfunction has been linked to a variety of diseases including neurodegenerative illnesses, metabolic disorders, lysosomal storage disorders and cancer [<u>3</u>].

Alpha one antitrypsin (A1AT) is a serum protease inhibitor primarily synthesized in the liver and serves as a pulmonary neutrophil elastase neutralizer to protect the lung from self-elastolytic digestion. Thus, it plays a critical role in maintaining proper pulmonary proteolytic-inhibitor balance[<u>6,7</u>]. A1AT is a highly polymorphic protein, over 120 mutations in the *SERPINA1* gene have been identified and most mutation caused disease pathogenesis manifests as pulmonary emphysema because the loss of function as elastase neutralizer. These mutated A1AT caused manifestation is called alpha one antitrypsin deficiency (AATD). One genetic mutation result for a mutant protein ATZ with a E342K substitution exert the most sever phenotype. ATZ has misfolded conformation and prone to aggregate into insoluble polymers and The abnormal ATZ polymerization could result in liver cirrhosis [<u>7,8</u>]. Lung diseases resulting from AATD are treated with intravenous infusions of A1AT from donated human plasma, but this therapy does not mitigate liver damage caused by hepatic ATZ aggregates [<u>9</u>].

It has been established that, the ubiquitin–proteasome and autophagy–lysosomal pathways represent the two major routes for ATZ’s bio-digestion [<u>10,11</u>]. Enhancing autophagy using chemical compounds have all been shown to prohibit A1AT protein accumulation and alleviating cell injury[<u>12</u>]. To date, there have been no studies exploring the role of CMA in the regulation of ATZ catabolism. Theoretically, proteins with KFERQ motif could be recognized by HSC70 and delivered for CMA degradation. Searching the sequence of ATZ, there is a typical pentapeptide motifs within the ATZ sequence and those amino acids are exposed on the surface of the protein molecular. This observation leads us to speculate that ATZ could be targeted for CMA, and manipulating CMA might offer a solution for ATZ digestion, preventing substantial polymerization.

## Material and methods

### Cell culture and plasmids transfection

HEK 293 and HepG2 cells were obtained from the National Collection of Authenticated Cell Cultures in Shanghai, China. The HeLa *ATG16L1-/-* and HeLa *BECLIN1-/-* cell lines were provided by Dr. Yue Xu at Shanghai Jiao Tong University, China. All cells were cultured in Dulbecco’s Modified Eagle Medium (DMEM, FI101-01, TransGen Biotech) supplemented with 10% fetal bovine serum (FBS, FI101-04, TransGen Biotech), 50 IU penicillin, and 50 µg/ml streptomycin under conditions of 37° C with a CO_2_ concentration of 5%. Following the manufacturer’s instructions for transfection reagents (Lip8000, Beyotime), cells at a confluence level of approximately 70% to 80% underwent plasmid transfection. Cells were then harvested for further analysis after 48 to 72 hours post-transfection.

### Antibodies and Chemicals

A1AT(8H10L18, ThermoFisher), LC3(AL221, Beyotime), Flag(AE005, AE169, ABclonal), ATG16L1(AE3637, ABclonal), BECLIN1(A21191, ABclonal), β-Actin(AC026, ABclonal), LAMP2A(AF1036, Beyotime), HRP Goat Anti-Rabbit IgG (H+L) (AS014, ABclonal), HRP Goat Anti-Rabbit IgG (H+L)(AS003, ABclonal), ABflo® 594-conjugated Goat Anti-Mouse IgG (H+L) (AS054, ABclonal), ABflo® 488-conjugated Goat Anti-Rabbit IgG (H+L) (AS053, ABclonal) Cycloheximide(01810, Sigma), DMSO(D8371, Solarbo), puromycin(P8230, Solarbo), streptomycin(P1400, Solarbo), AR7(HY-101106, MedChemExpress), QX77(HY-112483, MedChemExpress), MG-132(HY-13259, MedChemExpress), Chloroquine(HY-17589A, MedChemExpress).

### Molecular cloning and ShRNA knockdown

*LAMP2A* and *HSC70* were cloned into pCDH-CMV (72265, Addgene) by PCR amplifying the ORFs from cDNA templates of HSC70 (19514, Addgene) and LAMP2A (86146, Addgene), followed by digestion with BamHI and EcoRI restriction enzymes. The molecular cloning was performed using T4 DNA ligase and the resulting constructs were transformed into DH5α E. coli cells. Plasmid purification and extraction steps were carried out using the Endo Free Mini Plasmid Kit II (DP118, TIANGEN). ShRNA sequences are detailed in supplementary Figure 1, and the corresponding shRNA plasmids were transfected into cells using Lip8000 transfection reagent (C0533FT, Beyotime), following the manufacturer’s protocol.

### Cloning of lentiviral expression constructs

To develop GFP-ATZ and LAMP2A-expressing HEK 293 cells in a stable manner within the pCDH-CMV vector using lentivirus, the *LAMP2A* and *ATZ* genes were amplified by PCR, specifically targeting the ORFs from the cDNA templates of LAMP2A or SERPINA1. The resulting cDNA insert of pCDH-CMV was then sub-cloned into the lentiviral backbone plasmid pCHD-CMV-IRES-pur, thus constructing the pCDH-CMV-GFP-ATZ or pCDH-CMV-LAMP2A plasmids. To generate lentiviral particles expressing the pCDH-CMV-GFP-ATZ or pCDH-CMV-LAMP2A plasmids, the corresponding constructs were co-transfected with the VSV-G and PVSV-2 helper plasmids into HEK 293 cells. The transfected cells were cultured for 4 days to ensure sufficient expression time. Subsequently, the culture medium containing viral particles was collected, filtered to remove debris, and used to infect HEK 293 cells, inducing efficient expression of the target proteins within the host cells.

### Discovery and Mutations of the CMA Recognition Motif “KFERQ-like Pentapeptide”

To demonstrate the involvement of CMA recognition motif, the complete amino acid sequence of the *SERPINA1* gene was obtained by accessing the UniProt database (P01009). A search was performed using the web server “KFERQ finder V0.8” (https://rshine.einsteinmed.edu/) to identify pentapeptide sequences associated with the CMA recognition motif. Within the amino acid sequence of *SERPINA1* (A1AT), a canonical pentapeptide fragment, QELLR, was found at positions 121-125. To modify the physical properties of amino acids within the targeting motif, as described by Orenstein and Cuervo [<u>13</u>], we introduced mutations into the CMA recognition motif (5’-CAGGAACTCCTCCGT-3’) resulting in 5’-GCGGCACTCCTCCGT-3’. The mutated motif was then sub-cloned into a pCDH-CMV plasmid.

### Fractionation of Cytoplasmic Soluble Proteins and Aggregates

The cells were lysed using RIPA lysis buffer. After centrifugation at 13,000g for 10 minutes at 4°C, the supernatant was collected as the cytoplasmic soluble fraction, and the pellet was resuspended in lysis buffer containing an additional 2% SDS. The resuspended pellet was further homogenized by puncturing with a 28-gauge needle ten times.

### Immunoblotting

Cell lysates were prepared in RIPA buffer to extract total protein. The obtained lysates were then subjected to centrifugation at 17,000g for 20 minutes at 4°C to separate the soluble lysates from the insoluble microspheres. Protein concentrations were determined using the Bicinchoninic acid method, as directed by the manufacturer’s instructions using the Pierce BCA protein assay kit (23235, Thermo). Equal amounts of protein were transferred to a polyvinylidene fluoride (PVDF) membrane through sodium dodecyl sulfate polyacrylamide gel electrophoresis (SDS-PAGE). The primary antibody was added and incubated overnight at 4°C, followed by incubation with the secondary antibody at room temperature for 45 minutes. Blot was performed using a chemiluminescence kit (BeyoECL Plus P0018S, Beyotime) and visual detection was performed using a western blotting detection system (Tienong 5200, China). The gray values of protein bands in each group were subsequently quantified using ImageJ software, and the ratio of target bands to β-actin bands was utilized as the measure of protein expression level. The experiment was repeated three times with distinct protein samples.

### Immunoprecipitation

To obtain the whole cell extract, the cells were treated with lysis buffer containing protease inhibitor, and the resulting supernatant was collected. Magnetic beads (AC047, AE097, ABclonal) were pre-treated with antibody binding and subsequently mixed thoroughly with the extracted total cell protein samples. The mixture was then subjected to rotational agitation and incubated overnight at 4°C to facilitate the formation of immune complexes. The protein-bound magnetic beads were isolated using a magnetic separation rack, and the supernatant from the heated beads was analyzed through SDS-PAGE and western blot.

### Immunofluorescence

The cells were washed three times with pre-chilled PBS (P1020, Solarbio), for 1 minute each time. Subsequently, the cells were incubated at 37°C for 20 minutes in an immunostaining fixation solution (P0098, Beyotime) and then underwent three additional washes with PBS, each lasting 5 minutes. Following a 25-minute incubation at room temperature in an immunostaining permeabilization solution (Triton X-100, P0096, Beyotime), the cells were again washed three times with PBS for 5 minutes each. Next, the cells were permeabilized, and non-specific sites were blocked using an immunostaining blocking solution (P0102, Beyotime) at room temperature for 1 hour. The primary antibodies, diluted in an immunostaining primary antibody dilution solution (P0103, Beyotime), were then incubated overnight at 4°C. On the following day, the primary antibodies were removed, and a wash with immunostaining washing solution (P0106, Beyotime) was performed on the cells. Subsequently, the cells were incubated at room temperature for 1 hour with secondary antibodies, diluted in an immunostaining secondary antibody dilution solution (P0108, Beyotime). The cells were washed three times with PBS. Finally, the cells were mounted onto slides using a fluorescence decay mounting medium containing DAPI (S2110, Solarbio). Confocal microscopy images were captured, processed using a laser scanning confocal microscope (TCS SP5II, Leica), and analyzed using ImageJ.

### Cellular GFP-ATZ protein clearance rate assessed by flow cytometry

HEK 293 cells expressing GFP-ATZ were transfected with a plasmid for 48 hours. After centrifugation, the cells were resuspended in phosphate-buffered saline and analyzed using FACScan flow cytometry (BD LSR Fortessa X-20) for GFP fluorescence (488-nm laser excitation, 530/30 filter for detection). A total of 3×10^5^ cells underwent scanning to exclude cellular debris based on forward and side-angle scatters (FSC and SSC). Cells expressing GFP were gated, and the percentage depletion of GFP-positive cells was calculated relative to the control group. The clearance rate of GFP-ATZ protein is assessed by measuring the average fluorescence intensity. The clearance rate of GFP-ATZ protein is assessed by measuring the average fluorescence intensity.

### Cycloheximide Chase Analysis

The CHX chase experiments involved a 48-hour cell culture period following plasmid transfection. Protein synthesis was blocked using CHX treatment at a concentration of 100 μg/ml. Cells were collected at various time points, lysed with RIPA buffer, and subjected to SDS-PAGE and immunoblotting analysis with specific antibodies for protein detection.

### Real-time Quantitative RT-PCR Assay

Total RNA was extracted from GFP-ATZ expression HEK 293 stable cell line using TRIZOL (15596026, Thermo) according to the manufacturer’s instructions. A total of 1μg RNA was used for reverse transcription (QIAGEN, 205311). qRT-PCR was performed using gene-specific primers and SYBR Green Real-time PCR Master Mix (Toyobo, Osaka, Japan). The gene expression levels were normalized to β-actin. Real-time PCR and data collection were conducted on a Bio-Rad CFX96 Connect instrument. The primer sequences used for PCR were as follows: *LAMP2A* and *LAMP2B* and *LAMP2C* forward oligo: 5′-GAA GGA AGT GAA CAT CAG CATG-3′, *LAMP2A* reverse oligo: 5′-CTC GAG CTA AAA TTG CTC ATA TCC AGC-3′, *LAMP2B* reverse oligo: 5′-CAA GCC TGA AAG ACC AGC ACC-3′, *LAMP2C* reverse oligo: 5′-CTC GAG TTA CAC AGA CTG ATA ACC AGT AC-3′, *HSC70(HSPA8)* Forward oligo: 5′-CAC TTG GGT GGA GAA GAT TTTG-3′, *HSC70 (HSPA8)* reverse oligo: 5′-CTG ATG TCC TTC TTA TGC TTG C-3′, *SERPINA1* forward oligo: 5′-GGC TGA CAC TCA CGA TGAAA-3′, *SERPINA1* reverse oligo: 5′-GTG TCC CCG AAG TTG ACA GT-3′.

### Transcriptional analysis

The KFEE pathway and GO term enrichment were perform using the Hiplot Pro, a comprehensive web service for biomedical data analysis and visualization. It involved utilizing the clusterProfiler package (v.4.5.0) in R software (v.4.2.2), as described by Wu et al [<u>14</u>]. Hiplot Pro served as the platform for importing and analyzing the data, while clusterProfiler facilitated various bioinformatics analyses. The analysis was carried out by using R software (v.4.2.2) package clusterProfiler (v.4.5.0) through Hiplot Pro (https://hiplot.com.cn/). Enrichment items with corrected *p*<0.05 were considered significantly enriched.

### Mitochondrial condition measurements

Cellular mitochondrial membrane potential was assessed using the Mitochondrial Membrane Potential Assay Kit (Beyotime, C2001S) with the TMRE dye. The principle of this assay kit is based on the response of TMRE dye to changes in mitochondrial membrane potential within cells. TMRE, a mitochondria-specific fluorescent probe with red fluorescence, exhibits distinct fluorescence signals upon binding to ATP within the mitochondria [<u>15</u>]. Cells were washed three times with chilled PBS and then incubated with TMRE fluorescent dye at 37°C for 30 minutes. Fluorescence intensity was measured using fluorescence microscopy and flow cytometry, with an excitation wavelength of 550nm and an emission wavelength of 575nm.

### Cell Viability

To assess cell viability, the Cell Titer-Glo® luminescent cell viability assay kit was used. The amount of ATP inside the cells was measured to indicate cellular metabolic activity. The assay measures the luminescent signal produced by the conversion of ATP to adenosine monophosphate (AMP) by the enzyme luciferase[<u>16</u>]. The treated cells were mixed with the reagent and agitated to induce cell lysis. The mixture was then incubated at room temperature to stabilize the luminescent signal. The luminescence intensity was recorded using the BioTek Synergy Neo2 multimode microplate reader.

### Statistical analysis

The statistical analysis of the data was performed using GraphPad Prism 9 (GraphPad Software). The experimental data was presented as mean ± SEM. Comparisons between two groups were conducted by unpaired two-tailed Student’s t-test for parametric data, and comparisons between multiple groups were conducted by one-way ANOVA. Statistical significance was set at *p*<0.05.*; *p*<0.01.**; *p*<0.001.***; *p*<0.0001.****; NC, no significance.

## Results

### CMA influence homeostasis of wide type and Mutant α1-Antitrypsin Z

In CMA, its activity is primarily modulated by changes in the levels of LAMP2A at the lysosomal membrane[<u>3</u>]. To evaluate CMA’s influence on ATZ, a stable cell line was created to express GFP-ATZ, which was subsequently transfected with incremental quantities of the LAMP2A-encoding plasmid. This resulted in a dose-dependent diminution of GFP-ATZ protein levels corresponding to the augmented expression of exogenous LAMP2A (Figure 1A, B). Simultaneously, a reduction in the standard CMA substrate GAPDH was observed, suggestive of enhanced CMA activity (Figure 1B).

**Figure 1:**
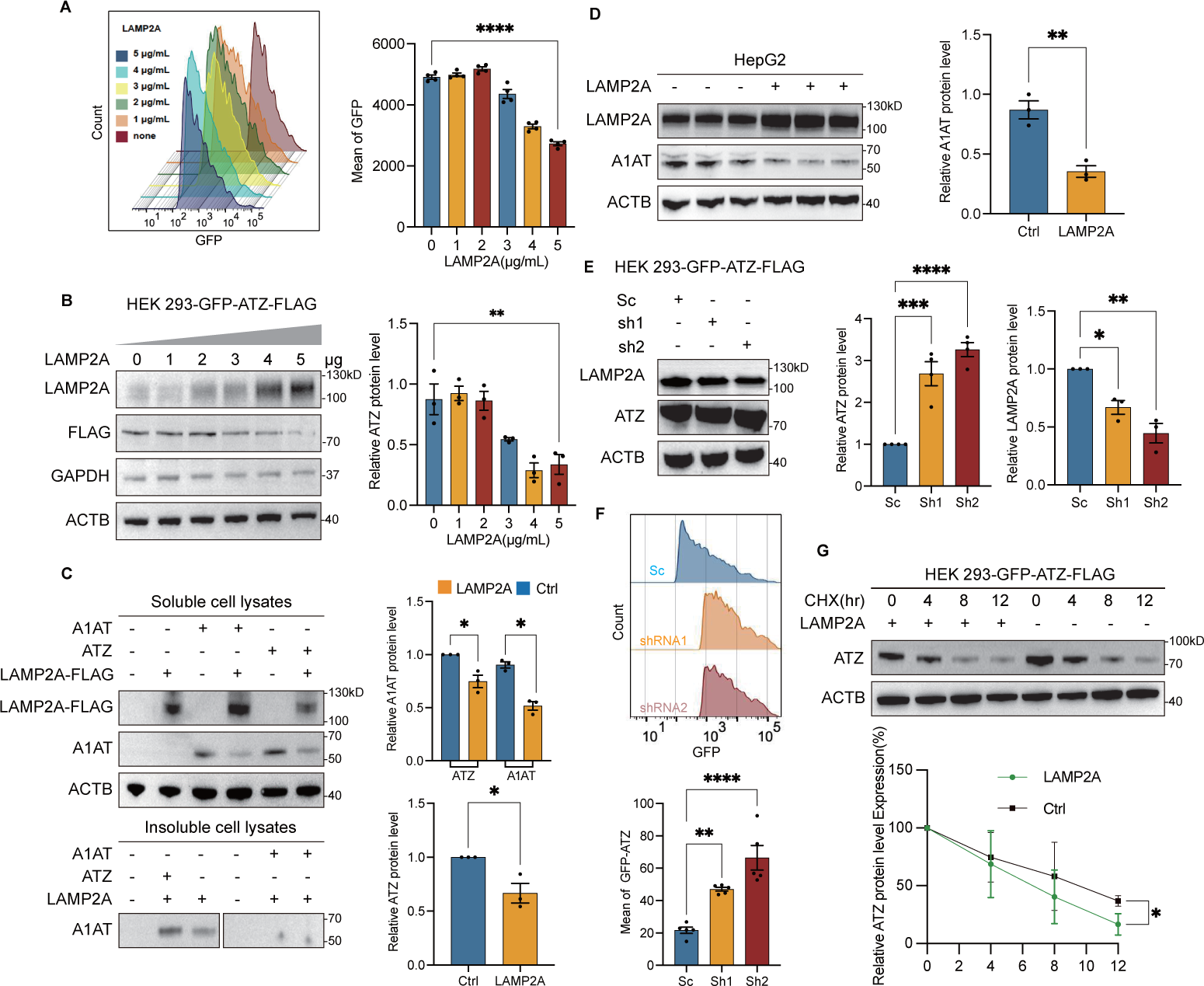
LAMPA2A influence mutant alpha one antitrypsin ATZ steady-state protein level. **A**: GFP-ATZ-expressing HEK 293 cells were transfected with increasing amounts (1, 2, 3, 4, or 5 μg/mL of cell medium) of a vector expressing LAMP2A for 48 hours. The fluorescence of GFP-ATZ was analyzed by flow cytometry. B: HEK 293 cells were transfected with increasing concentrations (1, 2, 3, 4, or 5 μg/mL of cell medium) of vectors expressing LAMP2A for 48 hours. The protein levels of GFP-ATZ relative to LAMP2A in cell lysates were quantitatively analyzed by measuring densitometry values from the blots. C: HEK 293 cells were co-transfected with LAMP2A and either ATZ or A1AT. After 48 hours, the cells were lysed using 0.1% Triton X-100 as a detergent, and the lysates were divided into Triton X-100 soluble cell lysates and Triton X-100 insoluble cell pellet fractions. Equal aliquots of cell lysates (30 μg) were loaded onto SDS-PAGE and immunoblotted with antibodies against ATZ and LAMP2A. Ctrl: pCDH empty plasmid. D: HepG2 cells were transfected with empty or LAMP2A-expressing vectors. After 48 hours of transfection, cell lysates were harvested and immunoblotted with the indicated antibodies. E: HEK 293 cells stably expressing GFP-ATZ were transfected with shRNA1 or shRNA2 to knock down LAMP2A protein expression for 48 hours. The levels of GFP-ATZ protein from different cells were quantitatively analyzed. Cells transfected with non-targeted scramble-shRNA were used as the control. Ctrl: scramble shRNA, sh1: LAMP2A-shRNA1, sh2: LAMP2A-shRNA2. F: HEK 293 cells stably expressing GFP-ATZ were transfected with shRNA1 or shRNA2 to knock down LAMP2A protein expression for 48 hours. The fluorescence of GFP-ATZ was analyzed by flow cytometry. G: GFP-ATZ-expressing HEK 293 cells were transfected with empty or LAMP2A-expressing vectors. Cells were treated with cycloheximide (CHX; 10 μg/mL) for the indicated times, starting 24 hours after transfection. The graph illustrates the quantitative changes in steady-state levels of GFP-ATZ relative to actin during the CHX treatment period. Data were presented as the mean ± SEM of 3 or more independent experiments. *p*<0.05.*; *p*<0.01.**; *p*<0.001.***; *p*<0.0001.****; NC, no significance. vs. indicated group.

Due to its aggregation-prone properties, the mutant protein ATZ leads to the formation of insoluble protein inclusions[<u>17</u>]. We thus resuspended the cell pellet in cell lysis buffer containing 2% SDS and dissect the CMA’s effect on insoluble ATZ. Unlike A1AT, we observed an aberrant amount of ATZ in the insoluble fraction. However, with the expression of LAMP2A, the insoluble ATZ level also significantly reduced (Figure 1C). Additionally, transfecting LAMP2A into the HepG2 hepatic cell line, which naturally expresses A1AT, resulted in a significant decline in endogenous A1AT levels upon LAMP2A overexpression (Figure 1D). This indicates that LAMP2A overexpression disrupts the homeostasis of both wild-type A1AT and ATZ.

To further validate the regulatory effect of LAMP2A on ATZ, we attempted to manipulate the endogenous levels of LAMP2A. We designed two short hairpin RNAs (shRNAs) that specifically target the cytosolic tail of the LAMP2A transcript, a region that differs between Lamp2 gene isoforms A and B, C (Supplementary Figure 1A, 1B). After transfecting the cells expressing GFP-ATZ with the two shRNA-coded vectors for 48 hours, a significant (>70%) reduction in LAMP2A mRNA levels was observed compared to cells treated with a scrambled shRNA, while no noticeable changes were detected in the transcripts of LAMP2B and LAMP2C (Supplementary Figure 1C,1D). Moreover, in the same samples, we detected a notable increase in GFP-ATZ protein levels (Figure 1E, 1F).

As the modulation of LAMP2A did not alter ATZ transcription (Supplementary Figure 2A), we speculate that the changes in ATZ levels may result from altered protein degradation dynamics. To confirm this, we blocked protein synthesis in the ATZ expression cells with cycloheximide (CHX) 24 hours after LAMP2A coding plasmids transfection and monitored the remaining ATZ at different time points post the drug treatment. The results indicated an accelerated degradation of ATZ in cells with LAMP2A overexpression, as evidenced by a decrease in half-life from 8 to 6 hours (Figure 1G). Additionally, overexpression of HSC70, another key component of CMA[<u>4</u>], also heightened the degradation of GFP-ATZ in a dose-dependent manner(Supplementary Figure 3A 、 B). These experiments collectively suggest that CMA plays a significant role in the proteostasis of both mutant ATZ and wild-type A1AT.

### ATZ is degraded through the CMA pathway mediated by LAMP2A

The above assays have provided evidences supporting CMA’s participation in regulating ATZ’s degradation, and we hypothesize that CMA directly targets ATZ for clearance. To corroborate this hypothesis, we conducted a pull-down assay of ATZ-flag protein and confirmed the association with LAMP2A (Figure 2A). Additionally, we stained endogenous LAMP2A and examined its co-localization with ATZ. We observed distinct CMA puncta, as indicated by LAMP2A, and a significant overlap between GFP-ATZ and LAMP2A (Figure 2B). We also examined the interaction between ATZ and HSC70 and found a potent binding of ATZ with HSC70 (Supplementary Figure 3C).

**Figure 2:**
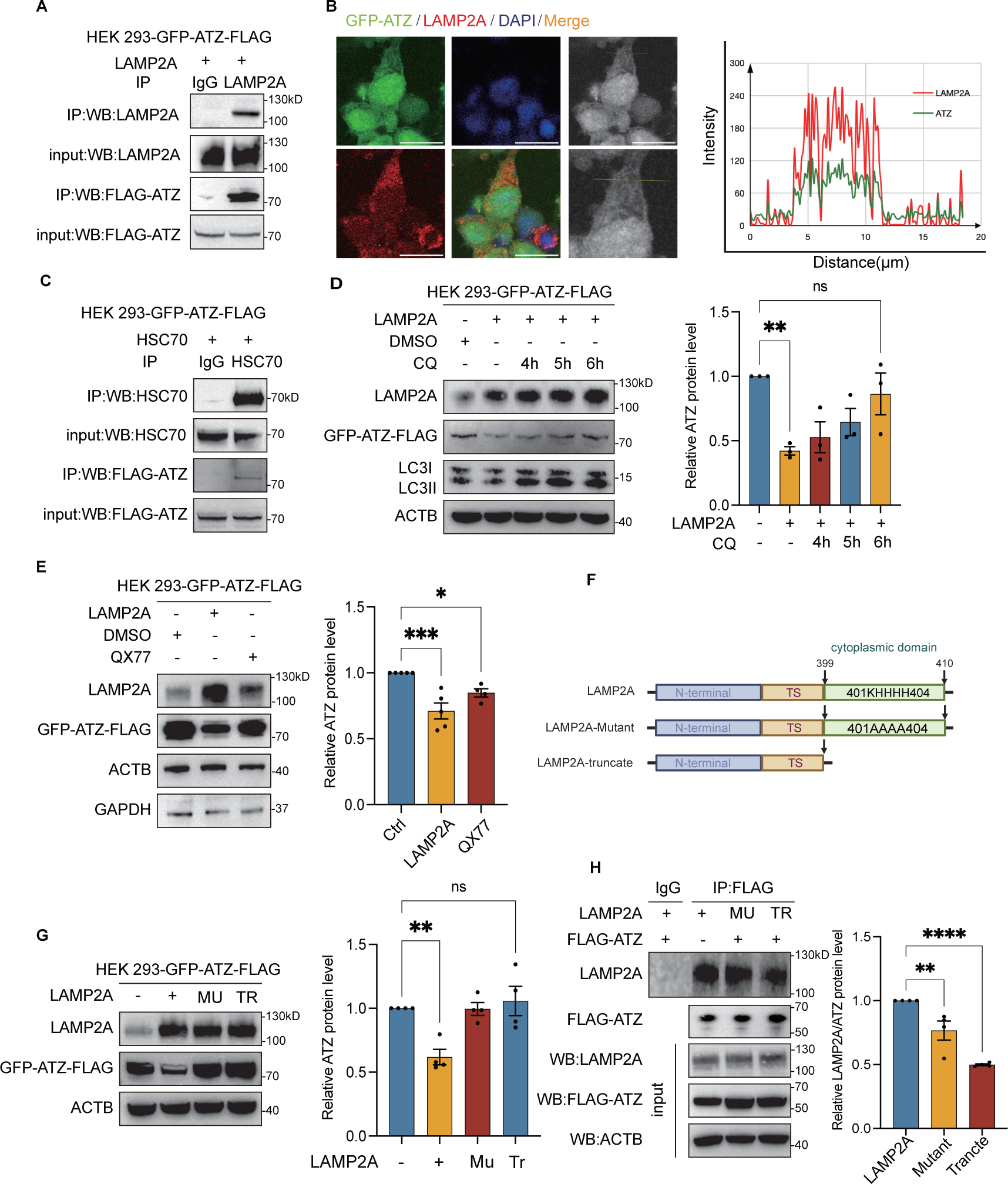
ATZ undergoes degradation via the CMA pathway facilitated by LAMP2A. A 、 C:HEK 293 stably expressing GFP-ATZ were transfected with an empty vector and LAMP2A or HSC70 for 48 hours, cells were lysed and subjected to immunoprecipitation with antibodies against IgG or Flag. Aliquots of IP proteins and input proteins were analyzed by immunoblot. B: HEK 293 stably expressing GFP-ATZ (green) were transiently transfected with HSC70 or LAMP2A for 48 h. Immunofluorescence microscopic images of GFP-ATZ (green) and LAMP2A(red) in HEK 293 stably expressing GFP-ATZ cells were obtained after staining with anti-LAMP2A antibodies. Each nucleus was stained with DAPI (blue). The co-localization of LAMP2A and ATZ-GFP were visualized as yellow color caused by superimposing of red and green. Plots of pixel intensity along the white line rows of images to the left of each plot, colors as in merged images. Scale bars: 10 μm D: GFP-ATZ-expressing HEK 293 cells were transfected with an empty vector or a vector encoding LAMP2A for 36 hours. Subsequently, they were treated with CQ (10μM) for an additional duration of 4, 5, or 6 hours. Following the treatment, the cells were lysed and underwent western blot analysis. E: HEK 293 cells expressing GFP-ATZ were treated with DMSO or QX77 (10μM) for 48 hours. Cell lysates were immunoblotted with the indicated antibodies. F: Schematic depiction of a full-length LAMP2A construct and LAMP2A-mutant construct. TS: Transmembrane structure. G: GFP-ATZ-expressing HEK 293 cells were transfected with full-length LAMP2A, LAMP2A-mutant, or LAMP2A-truncate constructs for 48 hours. The protein levels of GFP-ATZ in different transfections were quantitatively analyzed. TR: LAMP2A truncate, Mu: LAMP2A mutant. H: HEK 293 cells were transfected with LAMP2A, LAMP2A-mutant, or LAMP2A-truncate along with FLAG-ATZ for 24 hours. The cell lysates were then subjected to immunoprecipitation (IP) using an anti-Flag antibody, and the precipitates were analyzed by western blotting. TR: LAMP2A truncate, MU: LAMP2A mutant. Data were presented as the mean ± SEM of 3 or more independent experiments. *p*<0.05.*; *p*<0.01.**; *p*<0.001.***; *p*<0.0001.****; NC, no significance. vs. indicated group.

The degradation of substrates by CMA relies on the acidic environment within lysosomes [<u>18</u>]. To investigate whether ATZ degradation is dependent on the lysosome, we treated cells overexpressing LAMP2A with lysosomal inhibitor chloroquine (CQ) for variable times. We observed that the level of CQ blocked the autophagic-lysosomal pathway, leading to accumulated levels of ATZ (Figure 2D). These evidences thus prove the directed accessibility of CMA to ATZ and its mediation of degradation depending on lysosomal hydrolysis activity. Consistent with this, we found that QX77[<u>19</u>], a CMA-specific activator, was able to boost LAMP2A’s level and eliminate the typical CMA substrate GAPDH, as well as GFP-ATZ (Figure 2E).

The cytosolic tail of LAMP2A, comprising 12 amino acids, is essential for the recruitment of HSC70-substrate complexes to lysosomes, facilitating their interaction via positively charged residues [<u>20,21</u>]. To elucidate how LAMP2A recognizes ATZ, we engineered plasmids for two LAMP2A variants: one encoding a truncated version lacking the C-terminal tail (LAMP2A-truncate), and another encoding a mutant wherein positively charged amino acids at positions 401 to 404 (401KHHH404) were substituted with alanines (401AAAA404) (Figure 2F, Supplementary Figure 4). These plasmids were introduced into cells expressing GFP-ATZ. Our findings revealed that only the wild-type LAMP2A significantly reduced ATZ levels, whereas both the LAMP2A-truncate and LAMP2A-mutant variants were ineffective (Figure 2G). Additionally, immunoprecipitation assays with Flag-tagged ATZ demonstrated markedly reduced co-immunoprecipitation with the truncated and mutant forms of LAMP2A (Figure 2H). These data further affirm that CMA facilitates ATZ degradation in a LAMP2A-dependent manner, highlighting the critical role of the C-terminal cytosolic tail in mediating this interaction.

### The 121QELLR125 pentapeptide motif is crucial in CMA mediated ATZ degradation

The CMA pathway differs from autophagy in its distinct substrate preference. Substrates degraded through the CMA pathway typically contain the classical “KFERQ” pentapeptide sequence, which serves as a recognized site by HSC70[<u>3</u>]. Using the “KFERQ finder V0.8” web server (https://rshine.einsteinmed.edu/), we identified a typical KFERQ motif (121QELLR125) within the ATZ protein. To investigate the role of this specific motif, we engineered a mutant variant, AA-ATZ (121AALLR125) (Figure 3A) and evaluated its binding affinity to HSC70 and LAMP2A through a CO-IP assay. Remarkably, the mutant demonstrated a marked decrease in binding to both LAMP2A and HSC70 compared to the ATZ protein (Figure 3B, 3C). Immunofluorescence assays further substantiated the diminished interaction between the mutant ATZ protein and LAMP2A (Figure 3D).

**Figure 3:**
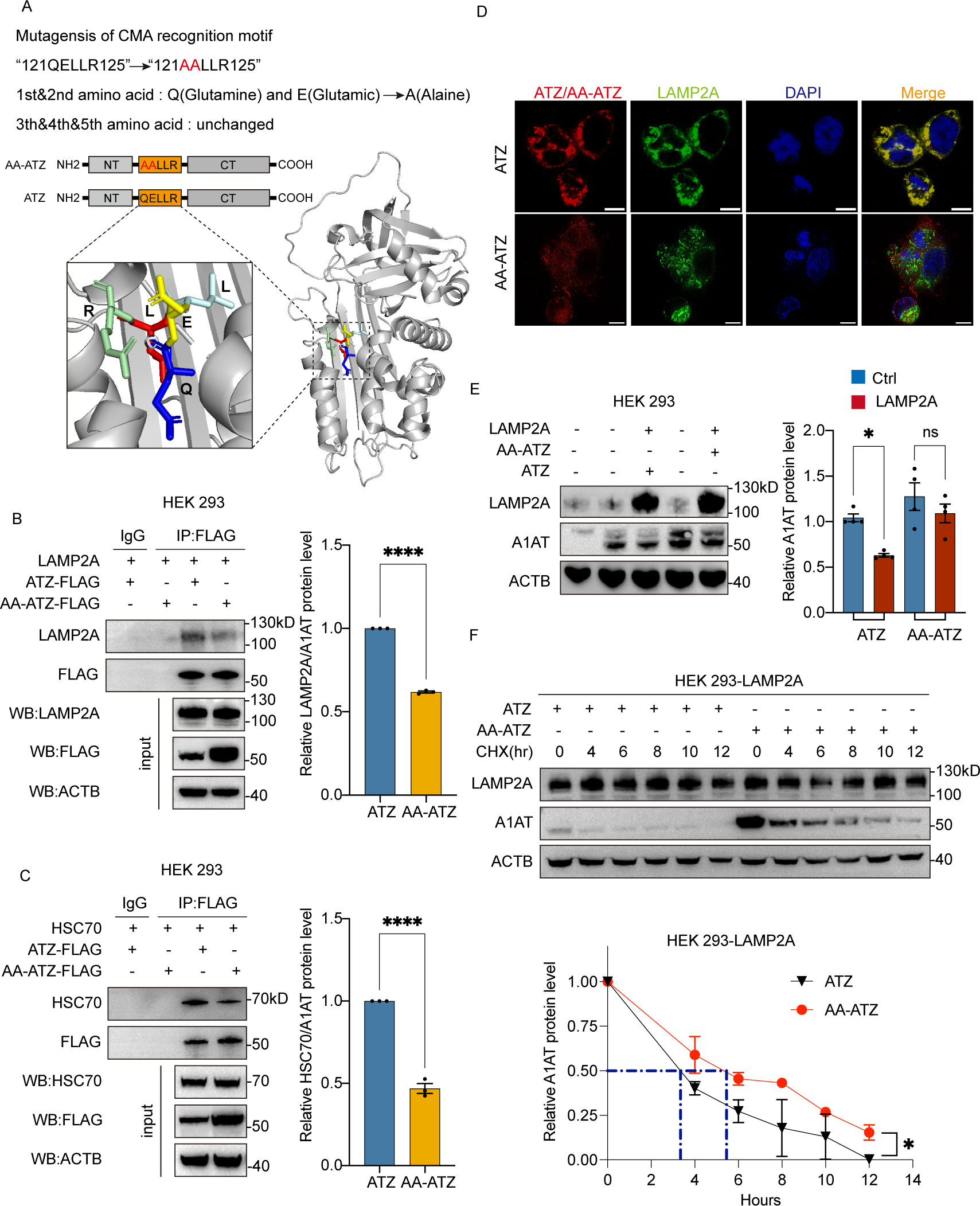
Pentapeptide 121QELLR125 on ATZ is critical in CMA mediated degradation. A: The cartoon illustrates the structure of alpha-1 antitrypsin, with the pentapeptide sequence (121QELLR125) highlighted in orange. The combined ribbon representation and stick model showing the overall structure and the CMA motif of ATZ protein. This model was generated using PyMOLTM based on the Protein Data Bank code 3CWM. B 、 C:HEK 293 cells transfected with LAMP2A or HSC70 and ATZ-FLAG or AA-ATZ-FLAG for 48h. Lysates were immunoprecipitated (IP) with either IgG, anti-LAMP2A antibody and probed for ATZ-FLAG, AA-ATZ-FLAG and LAMP2A or HSC70. D: HEK 293 cells were con-transfected for 48 hours with LAMP2A expressing vectors or vector coding wide type ATZ or AA-ATZ mutant. Representative confocal images showing the colocalization of ATZ-FLAG (red) or AA-ATZ-FLAG (red) and LAMP2A(green) in HEK 293T cells. Scale bars: 10 μm E: HEK 293 cells were co-transfected with LAMP2A expressing vectors or vectors encoding wild-type ATZ or AA-ATZ mutant for 48 hours. The cells were lysed, and western blot analysis was conducted to assess the expression of GFP-ATZ. F: HEK 293 stably expressing LAMP2A cells were transfected with wide type ATZ or AA-ATZ expressing vectors. The transfected cells were cultured for 24 h before being further incubated with cyclohexamide (CHX;10 μg/ml) for the indicated time. The levels of A1AT at different time points were detected by Western blot.

In addition, we observed that the basal protein level of AA-ATZ was significantly higher than that of ATZ under the condition of equal transfection of ATZ or AA-ATZ plasmids, regardless of whether LAMP2A was overexpressed or not (Figure 3E). We hypothesized that this discrepancy was caused by the inefficient degradation of mutant AA-ATZ by CMA. To prove this, we compared the degradation rate of ATZ with that of AA-ATZ under conditions of high CMA activity with a CHX chasing experiment. The results showed that, indeed, with the activation of CMA, the degradation rate of AA-ATZ was slower than ATZ, with a protein half-life reduced from 5.2 hours to 3.5 hours (Figure 3F). These results underscore the importance of the CMA pentapeptide 121QELLR125 in ATZ for its degradation through the CMA pathway. They further demonstrate that ATZ is *bona fide* substrate of CMA.

### CMA degrade ATZ independent of macroautophagy

With above assays, we confirmed that ATZ could be degraded via the CMA pathway. However, there are complicated compensatory regulatory mechanisms between macroautophagy and CMA [<u>22-24</u>]. To mitigate the influence of macroautophagy, we employed autophagy-deficient HeLa *ATG16L1-/-* cells[<u>25</u>] to reassess our findings. In wild-type HeLa cells, we observed that serum starvation not only activates CMA but also macroautophagy, as indicated by increased levels of LAMP2A and LC3II, alongside enhanced ATZ degradation (Figure 4A). Conversely, in HeLa *ATG16L1-/-* cells subjected to identical starvation conditions, the absence of LC3II formation confirmed the disruption of autophagy, yet ATZ protein levels still diminished (Figure 4B). Furthermore, in HeLa *ATG16L1*-/-cells, the overexpression of exogenous LAMP2A resulted in dose-dependent degradation of ATZ, which was similarly effective in wild-type alpha-1 antitrypsin A1AT and insoluble ATZ aggregates (Figures 4C, D). By employing CMA-specific activators, AR7 and QX77 [<u>26,27</u>], we noted identical degradation patterns for the prototypical CMA substrates GAPDH and ATZ (Figure 4E). Further investigations into the pentapeptide’s influence in autophagy-deficient cells showed that AA-ATZ exhibited a lower degradation efficiency than ATZ (Figure 4F). These data suggesting that CMA is capable of degrading ATZ independently of macroautophagy.

**Figure 4:**
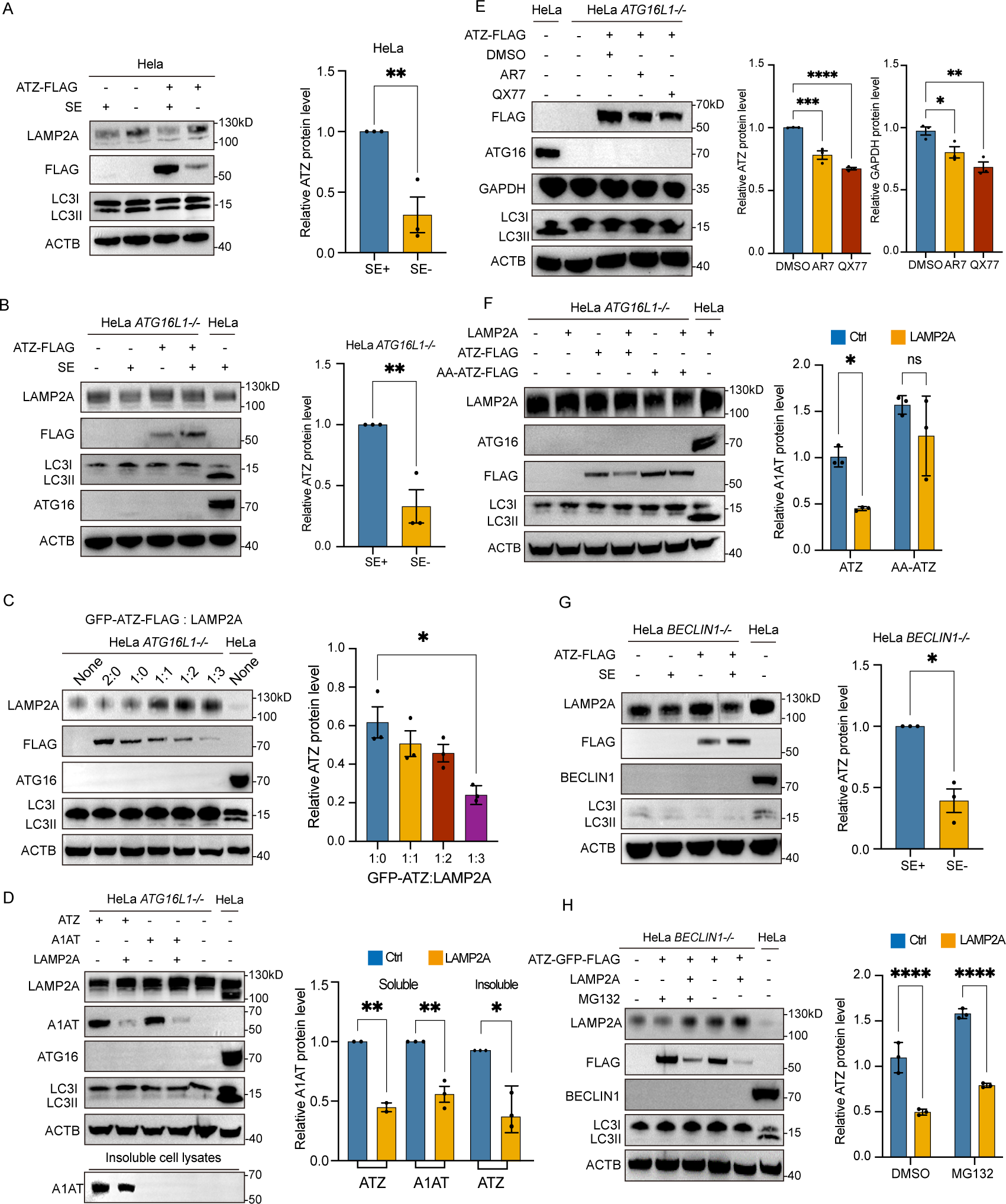
ATZ was a bona fide substrate of CMA. Ectopic expression of Lamp2A in macro-autophagy deficient cell causes reduction of ATZ. A: Hela *ATG16L1-/-* cells were transfected with ATZ-GFP-FLAG and amounts of a LAMP2A or expressing vector. Cells were lysed for Western analysis. B: HeLa *ATG16L1-/-* cells were transfected with LAMP2A and ATZ-FLAG or A1AT vectors. After 48 hours, cells were lysed using 0.1% Triton X-100 as a detergent, and the lysates were fractionated into Triton X-100 soluble cell lysates(n=2) and Triton X-100 insoluble cell pellet fractions(n=3). Equal amounts of cell lysates (30 g) were loaded onto SDS-PAGE and subjected to immunoblotting using antibodies against ATZ and LAMP2A. C: HeLa *ATG16L1-/-* transfected with FLAG-ATZ for 36h, treated with QX77(10μM) or AR7(10μM) for 24h, cell lysates were harvested and immunoblot with the indicated antibodies. D 、 E 、 F: Transfection of the ATZ-FLAG plasmid was performed in HeLa, HeLa *ATG16L1-/-*, and HeLa *BECLIN1-/-* cells. After 48 hours, the cells were subjected to free DMEM for 24 hours, followed by lysis of the cells for immunoblotting analysis. G: HeLa *BECLIN1-/-* cells were transfected with ATZ-FLAG and an Empty vector or LAMP2A for 36 hours and then treated with MG132 (20 μM) for an additional 4 hours. Cells were lysed and subjected to Western blot analysis. H: Immunoblotted assay of HeLa *ATG16L1-/-* cells transfected with LAMP2A and ATZ-FLAG or AA-ATZ-FLAG mutant. Data were presented as the mean ± SEM of 3 or more independent experiments. *p*<0.05.*; *p*<0.01.**; *p*<0.001.***; *p*<0.0001.****; NC, no significance. vs. indicated group.

In addition to its role in autophagy, *ATG16L1* has been implicated in anti-inflammatory responses and the regulation of hormone secretion[<u>28,29</u>]. To rule out the influence of *ATG16L1*’s non-autophagic functions, we utilized another autophagy deficient model, the HeLa *BECLIN 1-/-* cell line [<u>30,31</u>]. In the HeLa *BECLIN* 1-/-cells, a similar decrease in ATZ protein levels was observed after inducing CMA through serum starvation and LAMP2A overexpression (Figure 4G, H). Notably, this decline in ATZ persisted despite the inhibition of the proteasome pathway when LAMP2A was overexpressed (Figure 4H). These findings lend further support to the notion that CMA can independently facilitate the degradation of ATZ.

### CMA-mediated ATZ degradation mitigates cellular stress

Prior research indicates that ATZ aggregates leads to the cellular toxicity including mitochondrial damage[<u>32</u>]. We conducted a comparison of cell survival rates under normal conditions and after activating CMA. Overexpression of LAMP2A and HSC70 in cell lines stably expressing GFP-ATZ significantly improved cell viability (Figure 5A).Mitochondrial membrane potential alterations serve as a critical metric for mitochondrial damage assessment [<u>33</u>]. Using the TMRE probe to measure mitochondrial inner membrane potential[<u>15</u>], a notable increase in fluorescence intensity was detected upon overexpression of LAMP2A and HSC70, as compared to controls (Figure 5B,5C). This enhancement suggests a reestablishment of mitochondrial integrity, implicating the potential of CMA activation in mitigating mitochondrial damage.

**Figure 5:**
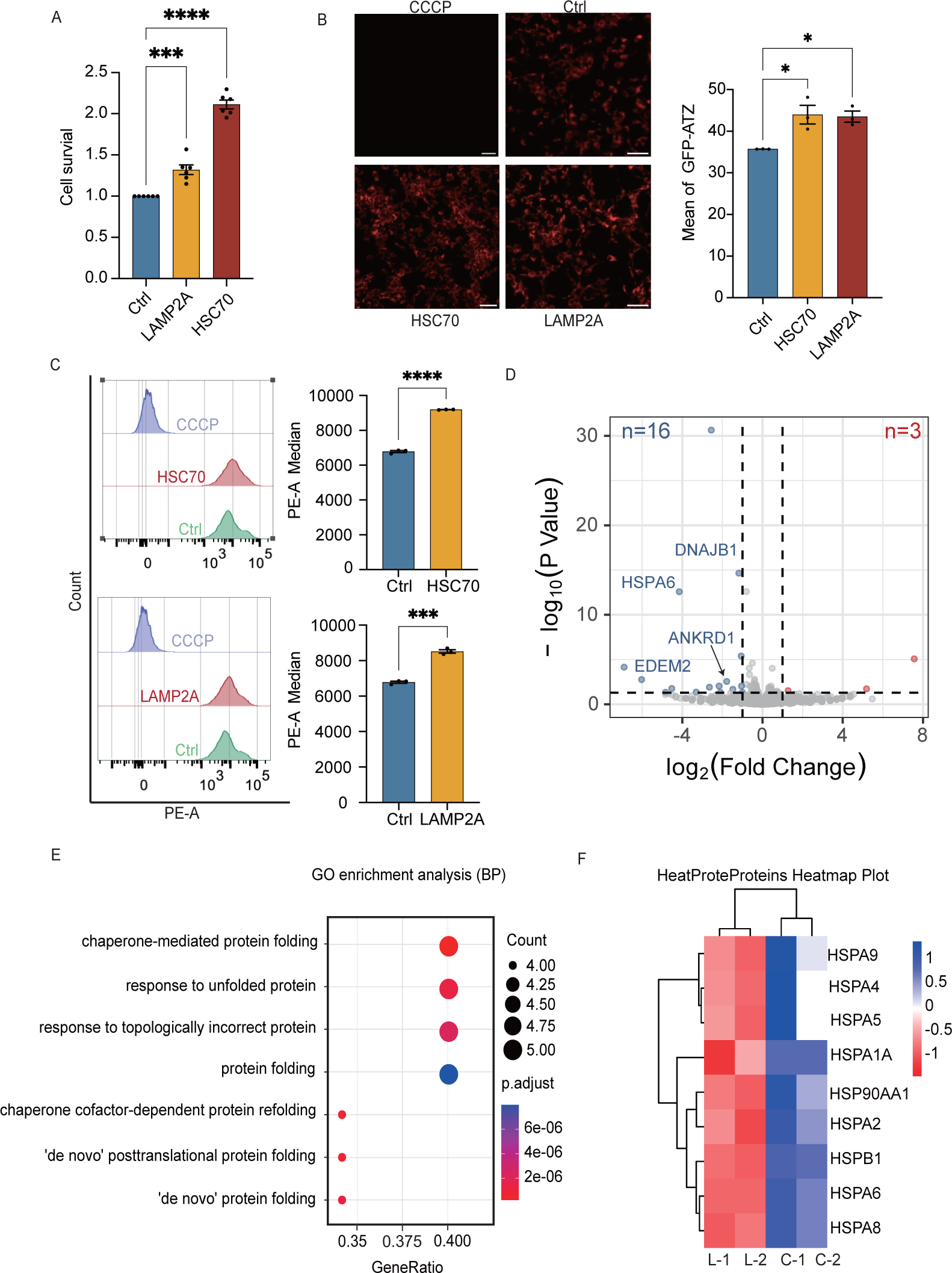
The degradation of ATZ facilitated by CMA alleviates cellular distress and ameliorates cellular states. A: HEK 293 cells expressing GFP-ATZ were transfected with LAMP2A or HSC70 for 48 hours. Cell viability analysis was performed using the CellTiter-Glo luminescent cell viability assay. B: TMRE signals in HEK 293 cells expressing GFP-ATZ were detected by fluorescence microscopy. Scale bars: 50 μm. CCCP: negative control. Ctrl: pCDH empty plasmid. C: HEK 293 cells expressing GFP-ATZ were transfected with either LAMP2A or HSC70, then stained with TMRE or CCCP for flow cytometry analysis. CCCP: negative control. Ctrl: pCDH empty plasmid. D: A volcano plot was generated to compare the log2 fold change (logFC) versus the p-value between control cells and LAMP2A-expressing cells. Grey circles represent RNAs with a p-value > 0.05, blue circles represent downregulated RNAs with a p-value < 0.05 and fold change greater than 2, while red circles represent upregulated RNAs with a p-value < 0.05 and fold change greater than 2, n=2. E: The expression levels of 16 genes displaying downregulation in HEK 293-GFP-ATZ cells under LAMP2A transfection conditions were subjected to Gene Ontology (GO) functional enrichment analysis (Biological Process). F: The corresponding heatmap highlights the disparity in RNA expression levels related to the heat shock protein family between the overexpression of the LAMP2A plasmid and the empty vector in GFP-ATZ-expressing HEK 293 cells.

Through transcriptome-wide analysis, we found that CMA activation via LAMP2A overexpression results in the upregulation of three genes and the downregulation of sixteen genes (Figure 5D, Supplementary Table). Notably, genes implicated in apoptosis and oxidative protein damage, such as DNAJB1, HSPA6, ANKRD1, and EDEM2, were markedly downregulated. Gene Ontology (GO) enrichment analysis further suggested that these downregulated genes are predominantly associated with chaperone-mediated protein folding (Figure 5E, Supplementary Figure 5). Additionally, KEGG pathway analysis underscored the downregulated genes’ roles in protein processing in the endoplasmic reticulum (ER) (Supplementary Figure 5). Considering the established function of heat shock proteins (HSPs) in the management of protein misfolding and ER stress, we performed a cluster analysis of HSP genes related to protein misfolding. The analysis revealed a decrease in the expression of various HSP genes upon CMA activation (Figure 5F), suggesting that enhanced ATZ degradation through CMA may alleviate ER stress and diminish the necessity for protein chaperones.

## Discussion

A comprehensive understanding of the catabolic mechanism responsible for the degradation of ATZ is essential for advancing therapeutic strategies for AATD. To date, the degradation of ATZ has been associated with both the proteasome and autophagy-lysosome pathways[<u>34</u>]. Multiple evidences indicated upregulated autophagic pathway in the AATD pathologic cells, and augmentation of autophagy has been demonstrated to facilitate the clearance of ATZ aggregates [<u>11,35</u>]. Furthermore, recent evidence has identified a noncanonical ER-to-lysosome-associated degradation pathway (ERLAD) that transports ATZ directly from the ER to the lysosome[<u>36</u>]. Despite this, the specific role of CMA in ATZ catabolism remains to be fully elucidated. This study presents biochemical evidence confirming CMA’s role in ATZ degradation and elucidates its underlying mechanisms.

CMA is a selective process where the chaperone protein HSC70 recognizes misfolded proteins, delivering them to LAMP2A for internalization. [<u>37</u>]. Our first piece of evidence supporting the involvement of CMA in the regulation of ATZ comes from the finding that LAMP2A directly interacts with ATZ and promotes its degradation (Figure 1 and 2). LAMP2A, produced by alternative splicing of the LAMP2 gene, is the only isoform implicated in CMA [<u>38,39</u>]. The selective silencing of LAMP2A leads to increased ATZ levels, underscoring CMA’s unique role in managing ATZ (Figure 1).

The structural distinctions between LAMP2 isoforms are localized to their C-terminal transmembrane region and cytoplasmic tail. The cytoplasmic tail’s positively charged residues are vital for substrate recognition in CMA [<u>39</u>]. Our research demonstrates that alterations to the cytoplasmic tail, either through deletion or modification of these key amino acids, compromise substrate binding and consequently hinder degradation (Figure 2G, H). This corroborates previous research underscoring the critical role of the cytoplasmic tail in the recognition of substrates by the CMA pathway.

CMA specific pentapeptide KFERQ motif in substrate is absolutely necessary for CMA mediated degradation, because it serves as a recognition site for HSC70 binding [<u>40</u>]. In this study, we identified a pentapeptide sequence, 121QELLR125, on ATZ and demonstrated that modifications to this sequence significantly weaken the interactions among ATZ, HSC70, and LAMP2A (Figure 3B, C). Variants of ATZ lacking this motif displayed a marked decrease in degradation rate, resulting in significant accumulation, even under normal physiological conditions (Figure 3E, F). The identification of 121QELLR125 motif further confirm ATZ as a bona fide substrate for CMA.

Macroautophagy and CMA, both lysosomal-dependent digestion pathways, are coordinately regulated in response to stress, with complex interplay and compensatory mechanisms. For example, the activation of CMA has been observed to suppress macroautophagy, while blocking CMA triggers an upregulation of macroautophagy [<u>22,24</u>]. Conversely, the induction of macroautophagy has been shown to diminish CMA activity [<u>23</u>]. In our study, we utilized two different autophagy-deficient cell lines to confirm that CMA acts independently to degrade ATZ (Figure 4). Conventionally, it is believed that in AATD affected cells, soluble ATZ is degraded by the proteasome system [<u>10</u>], whereas insoluble aggregates are cleared by autophagy-lysosome pathways and the ERLAD pathway[<u>11,35,36</u>]. The identification of CMA as a novel pathway for ATZ clearance brings us closer to achieving a comprehensive understanding of the catabolic processes involved in ATZ metabolism.

Microautophagy is another cellular degradation pathway conserved in mammalian cells, characterized by the direct engulfment of substrates by invagination of the lysosomal membrane[<u>41</u>]. The available data allow us to conclude that ATZ is a substrate of CMA, as it meets all the criteria for CMA substrates, including possessing the KFERQ-like pentapeptide, interacting with HSC70 and LAMP2A, and undergoing digestion within the lysosome. But it also raises another question: what is microautophagy’s role in ATZ’s catabolic regulation? While the precise involvement of microautophagy remains to be conclusively determined, emerging evidence suggests it could play a role in modulating ATZ levels. For example, overexpression of HSC70 exerts a greater effect on the clearance of ATZ and overall cellular recovery than LAMP2A overexpression does (Figure 5A). Such an effect is consistent with the process of endosomal microautophagy (e-MI), which selectively engulfs soluble proteins that present a KFERQ-like pattern, with HSC70 facilitation[<u>42</u>]. We postulate in that case overexpressed HSC70 may redirect a portion of ATZ to e-MI for degradation, offering additional cellular protection. Extensive research is necessary to thoroughly elucidating these mechanisms in this regard.

ER stress and mitochondrial damage are two typical characteristics of ATZ-induced cellular malfunctions[<u>43</u>,<u>44</u>]. In our study, we observed simultaneous alleviation of mitochondrial damage and restoration of cellular growth status upon the effective activation of CMA. Transcriptomic analysis revealed that CMA activation marginally altered global gene expression, with only 16 genes showing significant downregulation. Of these, four were intimately linked to protein folding, suggests a rebalancing of proteostasis due to the reduced presence of intracellular misfolded proteins [<u>45</u>]. The diminished expression of EDEM2, a gene implicated in endoplasmic reticulum-associated protein degradation [<u>46</u>], hints at the successful removing of misfolded proteins from the ER and a reduction in ER stress. Moreover, notable decreases in the expression of DNAJB1 and ANKRD1 both associated with apoptotic regulation[<u>47</u>] and ANKRD1[<u>48</u>], suggesting reduced cell apoptosis further confirming the positive role played by CMA towards maintaining cellular viability (Figure 5).

However, it is important to note that CMA serves as an essential intracellular protein degradation pathway. In theory, CMA targets proteins containing the KFERQ motif, which accounts for approximately 30% of soluble cytosolic proteins. Enhancing CMA activity would facilitate the removal of misfolded and oxidized proteins, and ultimately enhancing cellular viability[<u>22</u>]. Hence, in this context, Thus, the observed cellular recovery should not be exclusively credited to ATZ aggregate removal but also acknowledged as a broader impact of CMA in reducing general cellular stress by targeting various oxidized and misfolded proteins.

The aggregation of ATZ is a major contributor to the pathogenesis of AATD liver disease, prompting various attempts to halt or reverse the aggregation process and mitigate AATD progression. These strategies include inhibiting ATZ synthesis[<u>49-51</u>], enhancing secretion [<u>52,53</u>], blocking polymerization [<u>54,55</u>] and promoting degradation [<u>35,56</u>]. The identification of macroautophagy activation as a means to accelerate ATZ degradation has led to the investigation of autophagy-boosting drugs such as carbamazepine, rapamycin, and fluphenazine[<u>12</u>]. Our discovery of CMA involvement in the regulation of ATZ suggests that specific enhancers of CMA may offer new therapeutic options for AATD. Since the discovery of CMA pathway, there have been many attempts to discover small molecules for CMA manipulation. In recent years, a series of small molecular drugs that can activate CMA have been found. However, most CMA activators interfere with other cellular pathways, often causing disturbance in essential cellular processes, including biomaterial metabolism or protein synthesis, which hinders their application[<u>57</u>]. AR7 and QX77 both derivatives of retinoic acid, stands out as the most effective CMA inducer developed thus far [<u>19,27</u>].

In our model, we found the treatment with AR7 and QX77, enhancing CMA activity and clearing ATZ in both normal and autophagy-deficient cells (Figure1E and 4E). This finding suggests that CMA-specific activators could serve as potential small-molecule drug therapies for AATD. Previous studies conducted with Alzheimer’s disease mice have also demonstrated the clearance of pathological proteins when treated with AR7 derivatives, thereby providing support for their potential therapeutic implications [<u>58</u>]. Thus, performing future in vivo studies to investigate the impact of these CMA activators on ATZ clearance in animal models would hold significant value, allowing us to gain a deeper understanding of their therapeutic potential. In conclusion, our study establishes CMA as a previously neglected protein degradation pathway involved in regulating the catabolism of ATZ. As depicted in Figure 6, in addition to the proteasome and autophagy, ATZ can also undergo degradation via CMA. Specifically, the ATZ protein is recognized by HSC70 through the pentapeptide 121QELLR125 and delivered to LAMP2A to interact with its C terminal cytosolic tail on the lysosomal membrane, the LAMP2A then transporting the substrate into the lysosome for digestion. Activation of CMA, either through the overexpression of LAMP2A or the use of small molecular activators, enhances the digestion of ATZ, thereby slowing down its aggregation process and mitigating cellular stress.

**Figure 6:**
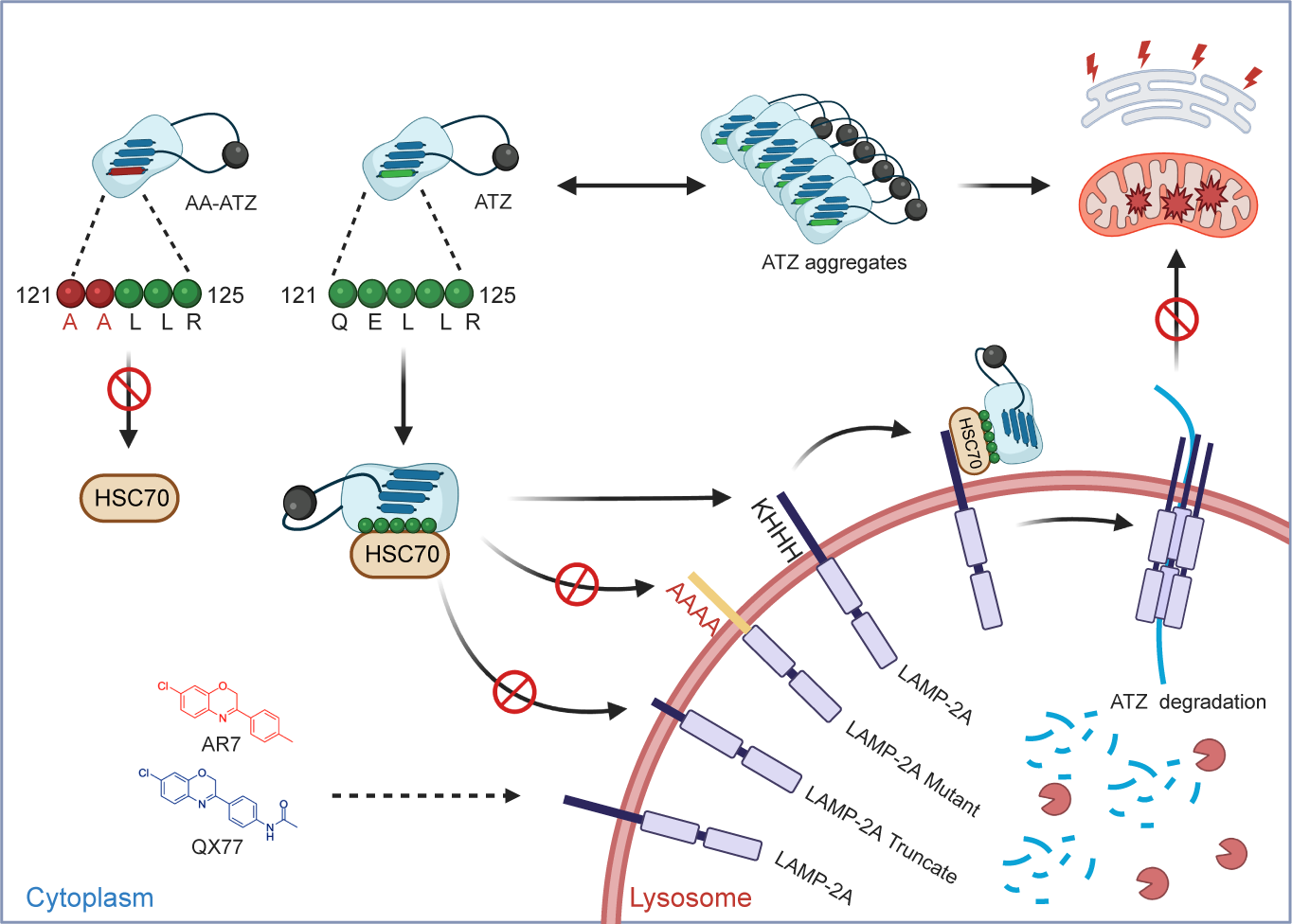
The schematic diagram illustrates the degradation mechanism of ATZ through the CMA pathway. Besides proteasome and autophagy pathways, CMA also plays a role in alpha-1 antitrypsin Z (ATZ) degradation. Specifically, the heat shock cognate 70 (HSC70) protein recognizes the ATZ protein through the pentapeptide sequence 121QELLR125 and transports it to lysosome-associated membrane protein 2A (LAMP2A) on the lysosomal membrane. Augmenting CMA activity through LAMP2A overexpression or AR7 utilization enhances ATZ digestion via this pathway, thereby alleviating cellular stress, including ER stress and mitochondrial impairment.

## Acknowledgments

This study was funded by: Tianjin Synthetic Biotechnology Innovation Capacity Improvement Project TSBICIP-CXRC-048; Public hospital reform and high-quality development demonstration project research fund, gastrointestinal tumors (2023SGGZ114); Major Project of Inner Mongolia Medical University (YKD2022ZD002).

## Author contributions

Hao Yang and Lang Rao contributed to the conception of the study; Jiayu Lin, Haorui Lu, Xinyue Wei, Yan Dai and Rihan Wu performed the experiment, Jiayu Lin, Hao Yang and Lang Rao did data analysis; Jiayu Lin, Hao Yang and Lang Rao prepared the manuscript.

