## Supplemental Figure1 for "Chaperone-mediated autophagy is an overlooked pathway for mutant α1-antitrypsin Z degradation"

### Supplementary Figure 1: Supplementary Figure 2: *LAMP2A* gene knockdown have no effect on mRNA levels of *LAMP2B* and *LAMP2C*

**A**

Scramble shRNA: CCTAAGGTTAAGTCGCCCTCG

**Forward oligo:**
5’ CACCGCCTAAGGTTAAGTCGCCCTCGCTCGAGCGAGGGCGACTTAACCTTAGGTTTTTG3’
**Reverse oligo:**
5’ AAACCAAAAACCTAAGGTTAAGTCGCCCTCGCTCGAGCGAGGGCGACTTAACCTTAGGC 3’

Human Lamp2A shRNA1:CTGCAACCTGATTGATTA

**Forward oligo:**
5’ CACCGCTGCAACCTGATTGATTACTCGAGTAATCAATCAGGTTGCAGTTTTTG3’
**Reverse oligo:**
5’ AAACCAAAAACTGCAACCTGATTGATTA CTCGAG TAATCAATCAGGTTGCAGC3’

Human Lamp2A shRNA2: GGCAGGAGTACTTATTCTAGT

**Forward oligo:**
5’ CACCGGGCAGGAGTACTTATTCTAGTCTCGAGACTAGAATAAGTACTCCTGCCTTTTTG3’
**Reverse oligo:**
5’ AAACCAAAAAGGCAGGAGTACTTATTCTAGTCTCGAGACTAGAATAAGTACTCCTGCCC3’

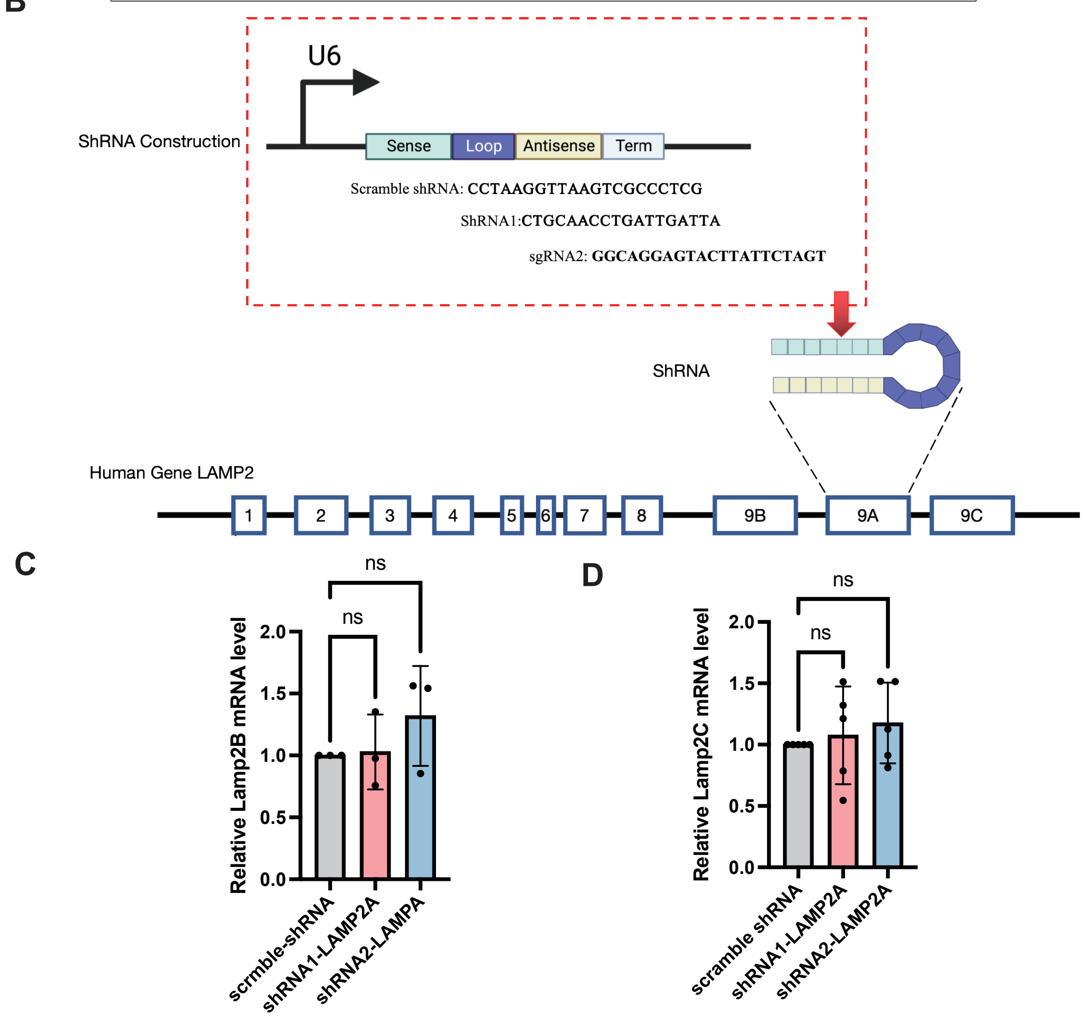


A:The DNA sequence for shRNA. B: Schematic graph of vector coding shRNA binding site of shRNA is on exon 9 of the LAMP2 gene.C、D: HEK293 cells stably expressing GFP-ATZ were transfected with shRNA1 or shRNA2 to knock down LAMP2A protein expression for 48 hours, after which RNA was isolated, and RT-PCR was performed for *LAMP2B* and *LAMP2C*.
