## Supplemental Figure2 for "Chaperone-mediated autophagy is an overlooked pathway for mutant α1-antitrypsin Z degradation"

Supplementary Figure 2: *LAMP2A* gene overexpression and knockdown have no effect on mRNA levels of ATZ

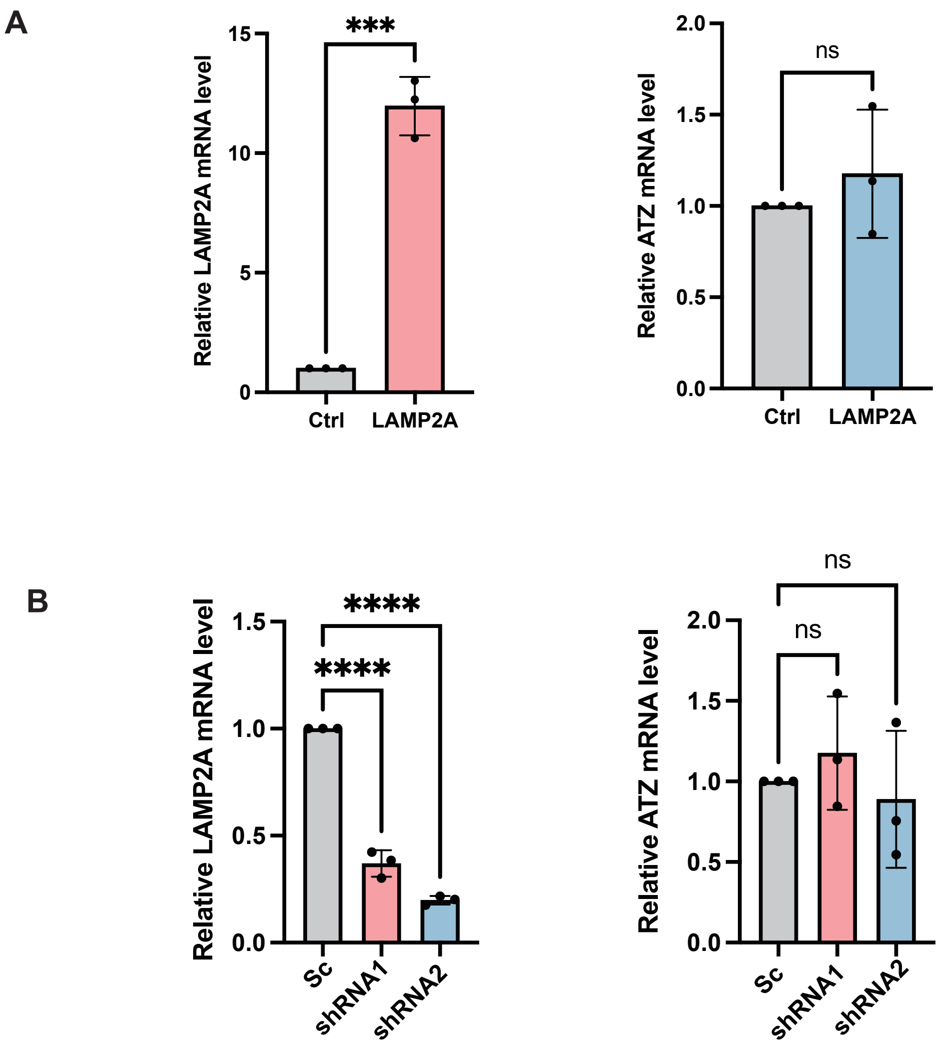


A: HEK 293 cells expressing GFP-ATZ were transfected with LAMP2A for 48 hours, after which RNA was isolated, and RT-PCR was performed for LAMP2A and ATZ.
B: HEK 293 cells were co-transfected with ShRNA1 or ShRNA2 and ATZ for 48 hours, after which RNA was isolated, and RT-PCR was performed for LAMP2A and ATZ.
Sc:scramble-shRNA, shRNA1:shRNA1-LAMP2A, shRNA2:shRNA2-LAMP2A
