## Supplemental Figure3 for "Chaperone-mediated autophagy is an overlooked pathway for mutant α1-antitrypsin Z degradation"

### Supplementary Figure 3: HSC70 regulates the protein levels of GFP-ATZ

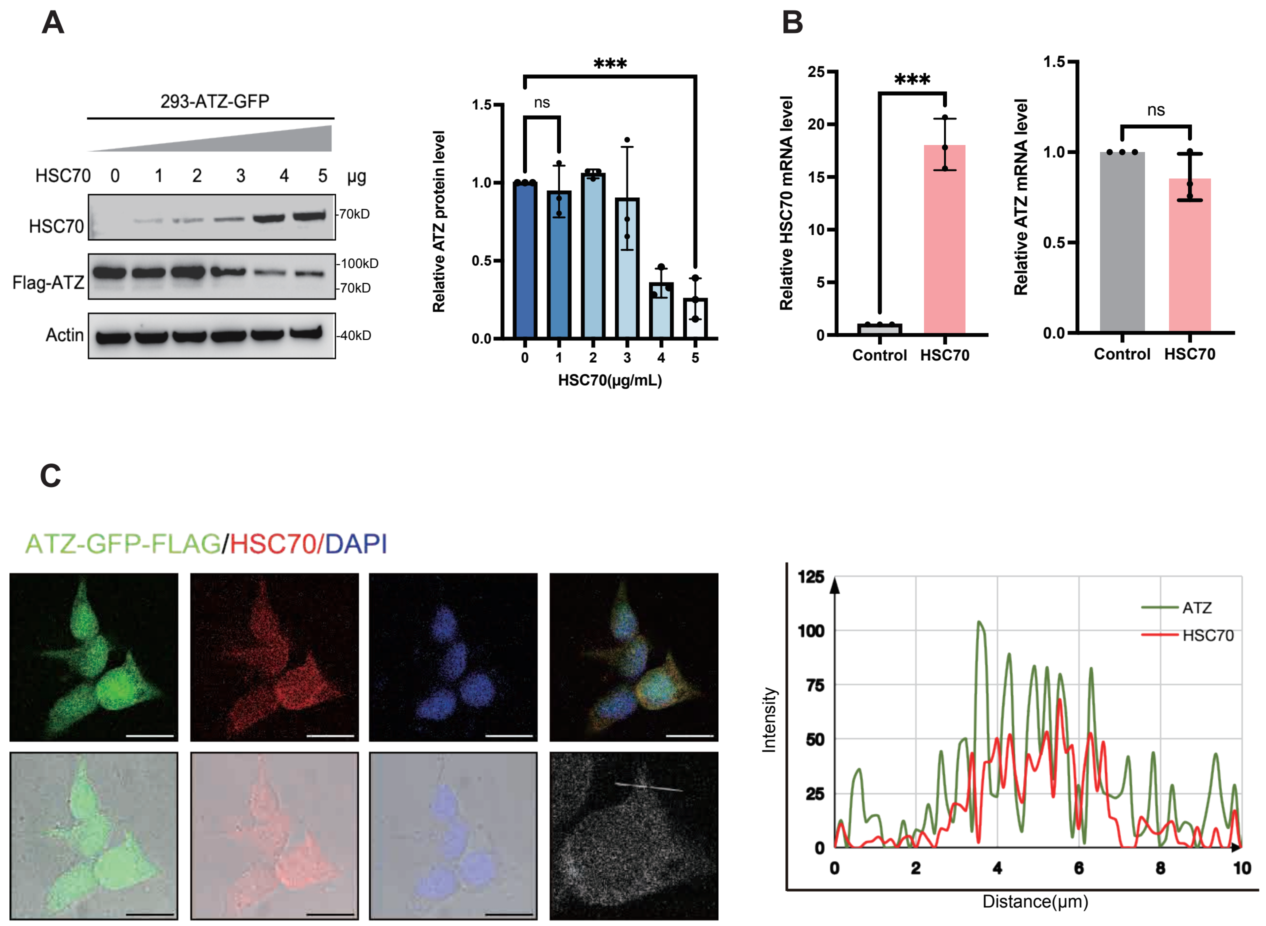


A: GFP-ATZ-expressing HEK 293 cells were transfected with increasing amounts (1, 2, 3, 4, or 5 μg/ml of cell medium) of a vector expressing LAMP2A. After transfection, the cells were lysed for Western blot analysis.
B: HEK 293 cells expressing GFP-ATZ were transfected with HSC70 for 48 hours, after which RNA was isolated, and RT-PCR was performed for LAMP2A and ATZ.
C:HEK 293 stably expressing GFP-ATZ (green) were transiently transfected with HSC70 for 48 h. Immunofluorescence microscopic images of GFP-ATZ (green) and HSC70 (red) in HEK 293 stably expressing GFP-ATZ cells were obtained after staining with anti-HSC70 antibodies. Each nucleus was stained with DAPI (blue). The co-localization of HSC70 and GFP-ATZ were visualized as yellow color caused by superimposing of red and green. Plots of pixel intensity along the white line rows of images to the left of each plot, colors as in merged images. Scale bars: 10 μm
