## Supplemental Figure4 for "Chaperone-mediated autophagy is an overlooked pathway for mutant α1-antitrypsin Z degradation"

Supplementary Figure 4:Construction of LAMP2A mutants and C-terminal cytoplasmic truncations

**A**

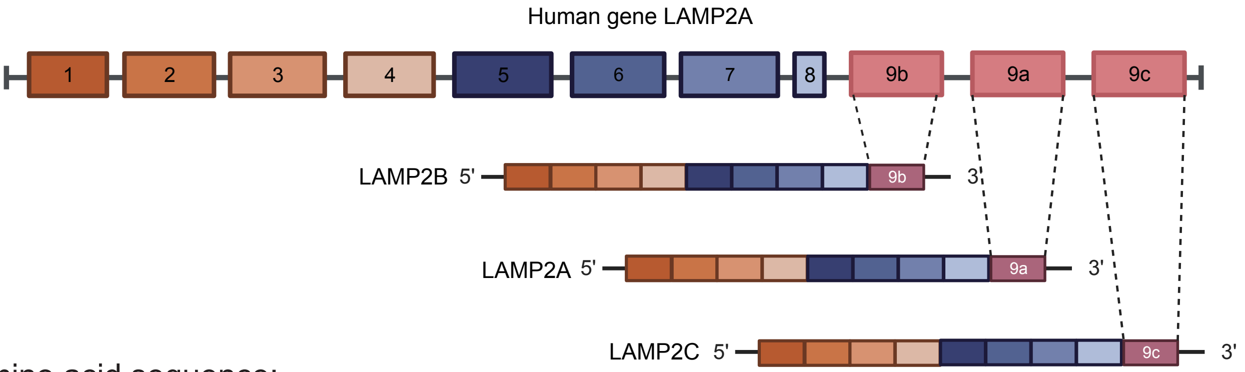


C-termina amino acid sequence:

*LAMP2A*: 375FLVPIAVGAALAGVLILVLLAYFIGLKHHHAGYEQF 410

*LAMP2B*: 375TILIPIIVGAGLSGLIIVIVIAYVIGRRKSYAGYQTL 410

*LAMP2C*: 3745LIPVAVGVALGFLIIVVFISYMIGRRKSRTGYQSV 411

**B**

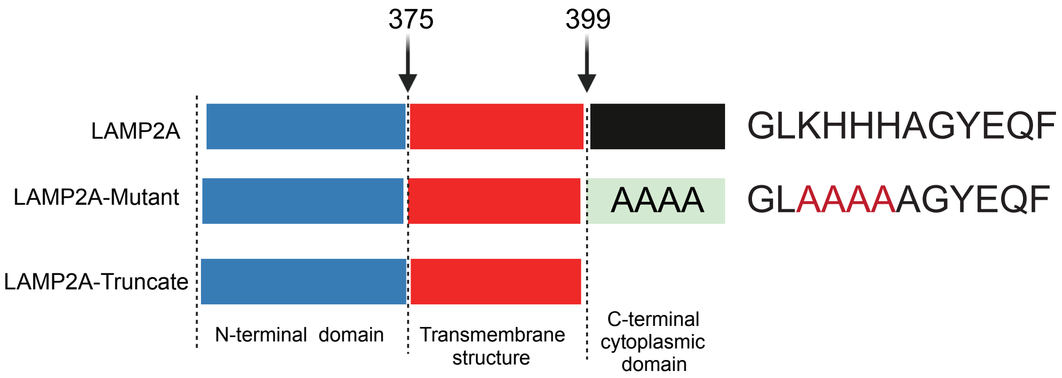


A: Alternative splicing of its pre-mRNA generates three isoforms: LAMP2A, LAMP2B, and LAMP2C, which share an identical lumenal domain (amino acids 1 to 374) at the N-terminus but differ in their transmembrane region (amino acids 375 to 398) and C-terminal cytoplasmic domain. Schematic depiction of LAMP2A and LAMP2B and LAMP2C construct.

B: Schematic depiction of a full-length LAMP2A construct and LAMP2A-mutant construct.
