## Supplemental Figure5 for "Chaperone-mediated autophagy is an overlooked pathway for mutant α1-antitrypsin Z degradation"

### Supplementary Figure 5: Downregulated genes in GO enrichment analysis and KEGG enrichment analysis

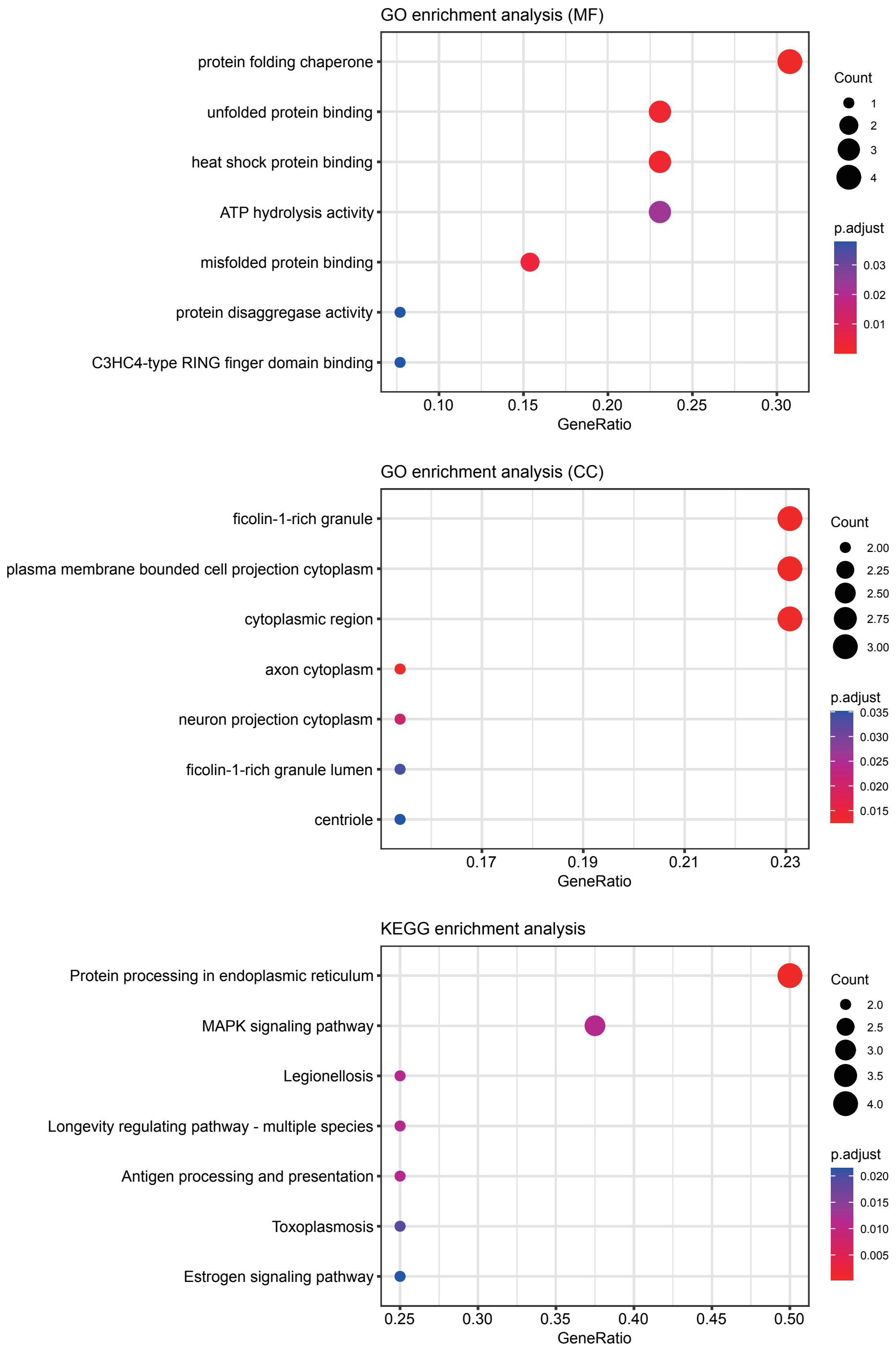


The 16 genes displaying downregulated expression in the HEK 293-GFP-ATZ cells under LAMP2A transfection conditions were subjected to Gene Ontology (GO) functional enrichment analysis (MF： Molecular function， CC:cellular component) and KEGG enrichment analysis.
