## Supplemental Table for "Chaperone-mediated autophagy is an overlooked pathway for mutant α1-antitrypsin Z degradation"

Supplementary Table: mRNA significantly down regulated by LAMP2A

| Other Gene ID | log2(LAMP2A/Control) | Pvalue(Control-vs-LAMP2A) |
|---|---|---|
| LAMP2 | 3.567725762 | 1.7069E-192 |
| DNAH17 | -2.554607904 | 2.29054E-31 |
| DNAJB1 | -1.178681508 | 2.22087E-15 |
| HSPA6 | -4.150401389 | 2.62317E-13 |
| DHRS2 | -1.06086095 | 4.0345E-06 |
| ZBED6CL | 7.563867925 | 8.35944E-06 |
| TMEM140 | -6.902650261 | 7.10599E-05 |
| TNFSF13 | -6.028888179 | 0.001713486 |
| ANKRD1 | -1.785434668 | 0.002780965 |
| ALPK3 | -1.044540164 | 0.009058812 |
| MMP24-AS1-EDEM2 | -2.167441329 | 0.009069338 |
| LOC112268437 | -2.647905394 | 0.012277967 |
| HOXA2 | -4.52528057 | 0.017226451 |
| CTAGE4 | 5.185064636 | 0.018649877 |
| GPNMB | -1.477786012 | 0.021170552 |
| MICOS10-NBL1 | 1.275346177 | 0.029887706 |
| HP | -3.306984987 | 0.043645593 |
| FCMR | -4.830798658 | 0.045203962 |
| RBM14-RBM4 | -1.004497171 | 0.046212602 |
| PPEF2 | -2.10525956 | 0.047062565 |

Significant differential transcriptional levels of genes were observed in the stable GFP-ATZ expressing cell line under conditions of transfection with LAMP2A plasmid, compared to transfection with empty vector. There were three upregulated genes and sixteen downregulated genes.
